# Near-atomic structures of human astrovirus capsids reveal a subunit-specific maturation mechanism

**DOI:** 10.64898/2026.06.22.733877

**Authors:** Kentaro Hiraka, Raymond N. Burton-Smith, Chihong Song, Kana Miyamoto, Kei Haga, Reiko Takai-Todaka, Akira Nakanishi, Kazuhiko Katayama, Kazuyoshi Murata

## Abstract

Human astrovirus (HAstV) is a major cause of acute gastroenteritis, yet the structural basis of capsid maturation and infectivity remains poorly understood. Here, we report near-atomic cryo-electron microscopy structures of HAstV serotype 4 in immature and mature states at 1.79 Å and 1.73 Å resolution, respectively. These structures reveal a subunit-specific maturation mechanism in which differential protease accessibility drives selective removal of spike (P2) domains. Structural comparisons show that only A-B dimers form a broad cavity beneath the P2 domains that is permissive for trypsin entry, whereas the corresponding region in C-C dimers is sterically restricted. Proteolytic maturation induces pronounced conformational rearrangements, particularly in subunit B, linking capsid geometry to asymmetric protease susceptibility. Despite these changes, the inner capsid architecture and presumptive RNA-associated densities remain conserved, indicating that genome release occurs at a later stage of infection. Together, our findings establish a mechanistic framework in which capsid geometry governs proteolysis-driven activation.

## Introduction

Astroviruses (family *Astroviridae*) are major enteric pathogens that infect a wide range of mammalian and avian hosts, representing a leading cause of acute gastroenteritis and diarrhea in young children worldwide ^1–4^. Despite their global prevalence and clinical impact, no approved vaccines or antiviral therapies are currently available, highlighting the urgent need to elucidate their infection and maturation mechanisms at the molecular level ^3–6^.

Human astrovirus (HAstV) is a non-enveloped, positive-sense RNA virus with an approximately 7.0-kb genome containing three open reading frames (ORFs) ^7^. ORF2 encodes the capsid protein (CP), initially synthesized as an 87-90 kDa precursor, termed VP90 (Fig. 1a) ^8,9^. Prior to viral release, the C-terminal acidic tail domains of VP90 are proteolytically cleaved by cellular proteases inside the infected cell, generating a 70 kDa core protein, termed VP70 ^10,11^. This VP70 product then assembles into T=3 icosahedral, non-infectious immature virus particles ^12^. The CP within these immature particles is structurally modular, comprising a conserved N-terminal capsid core [amino acids (aa) 1-415; containing a basic (Basic) domain (aa 1-72), a shell (S) domain (aa 73-262), and an outer core domain termed protruding 1 (P1) domain (aa 263-415)] and a protruding 2 (P2) domain (aa 416-647) ^13^.

**Figure 1.**
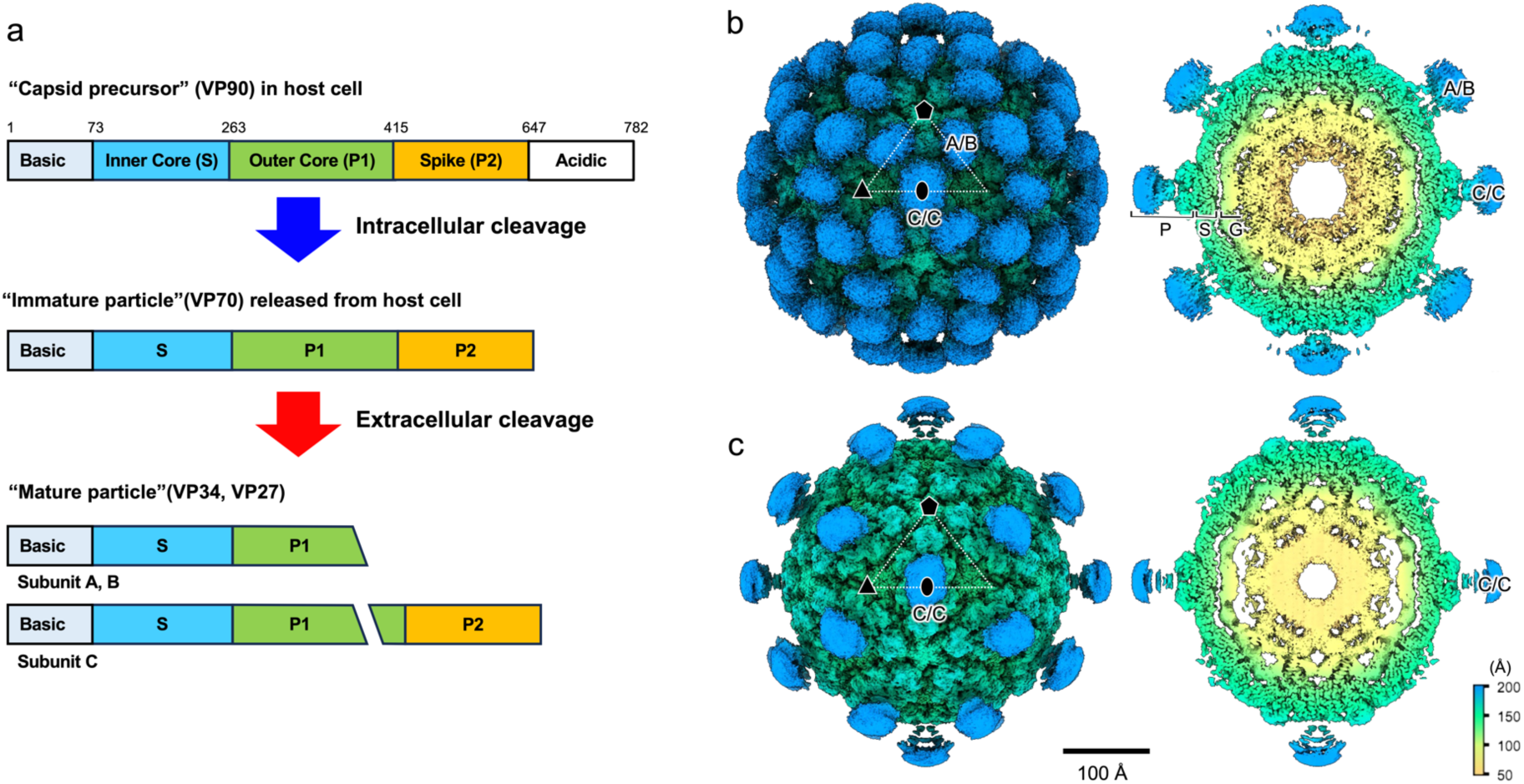
Maturation scheme and cryo-EM structures of immature and mature HAstV-4 particles. (a) Schematic representation of HAstV capsid protein processing. The capsid precursor VP90 (771 amino acids) comprises five domains: Basic (residues 1-70), S (71-263), P1 (264-429), P2 (430-646) and acidic (647-771). The acidic domain is cleaved intracellularly to generate immature VP70 particles, which are subsequently released from infected cells. Subsequent extracellular proteolysis produces VP34 and VP27/VP25, yielding mature particles. (b) Cryo-EM reconstruction of the immature particle shown as a surface view along the icosahedral twofold axis (left) and a central slice (right). (c) Cryo-EM reconstruction of the mature particle shown as a surface view along the icosahedral twofold axis (left) and a central slice (right). Maps are displayed with a Gaussian filter (1 Å) at a contour level of 0.5 σ. Radial distances are colour-coded as indicated. The P2 domains, capsid core (P1, S and Basic domains) and internal density are shown in blue, green and yellow, respectively. Icosahedral 2-fold, 3-fold, and 5-fold symmetry axes are indicated by oval, triangle and pentagon symbols.

To acquire infectivity, these extracellular immature VP70 particles must undergo robust proteolytic maturation driven by host tryptic cleavage ^14,15^. Biochemical models indicate that extracellular trypsin processes the VP70 precursor into distinct smaller fragments termed VP34, VP27, and VP25. During this transition, the VP34 and VP27 fragments remain tightly associated with the mature virion, whereas the P2-containing VP25 fragment is selectively released from the capsid shell ^16–18^. However, how the quasi-equivalent T=3 capsid geometry dictates this heterogeneous proteolytic processing and selective domain retention has remained fundamentally unresolved.

Here, we determine near-atomic cryo-electron microscopy structures of HAstV serotype 4 (HAstV-4) in both immature and mature states. These structures reveal that the quasi-equivalent capsid architecture dictates differential protease accessibility to the P2 domains. Our structural comparison demonstrates that variations between quasi-equivalent subunits correlate with trypsin susceptibility, clarifying the mechanism of heterogeneous proteolytic processing. Furthermore, we identify maturation-associated conformational changes throughout the capsid shell, along with unassigned internal densities distributed at roughly regular intervals that resemble the spacing of genomic RNA. Together, these findings characterize the structural transition of HAstV capsids, establishing a definitive framework to understand the subunit-specific maturation mechanism of the virus.

## Results

### Cryo-EM structures of immature and mature HAstV-4 particles

To define the structural basis of astrovirus maturation, the entire structures of immature and mature HAstV-4 particles were examined by cryo-electron microscopy (cryo-EM) (Fig. 1b,c). Single-particle analysis yielded reconstructions at global resolutions of 1.79 Å for the immature particle and 1.73 Å for the mature particle, respectively, enabling precise atomic modelling of the capsid core (Supplementary Tables 2 and 3 and Supplementary Fig. 1d,h).

Both immature and mature virions adopt a T=3 icosahedral architecture with an approximate outer diameter of 33 nm (Fig. 1b,c). Each asymmetric unit contains three quasi-equivalent subunits, designated A, B and C, which assemble into A-B dimers between the icosahedral 3-fold and 5-fold axes and C-C dimers along the strict icosahedral 2-fold axes. Although the overall resolution exceeded 2 Å, the local resolution varied across capsid regions, ranging from a highly ordered 1.4 Å within the inner core shell to >3.8 Å at the solvent-exposed periphery (Supplementary Fig. 1c,g). The inner capsid shell, comprising the N-terminal Basic, shell (S) and lower protruding (P1) domains, was resolved at near-atomic detail, allowing visualization of individual amino acid residues and their side chains (Supplementary Fig. 2). In contrast, the dimeric spike-forming upper protruding (P2) domains showed lower local resolution (Supplementary Fig. 1c,g), indicating significant intrinsic flexibility relative to the rigid icosahedral framework. In addition, concentric, diffuse internal densities were observed in both particle states, suggesting that the packaged viral single-stranded RNA genome does not adopt strict icosahedral symmetry.

### Atomic structures of capsid subunits A, B and C

Atomic models for the rigid capsid core region (residues 58-418) of subunits A, B and C were independently constructed for both particle states, excluding the flexible P2 domains (Fig. 2, Supplementary Fig. 2 and Supplementary Tables 2 and 3). In the immature particle, 350, 342, and 349 residues were successfully modeled for subunits A, B, and C, respectively, with Ramachandran outliers of 0.49%. In the mature particle, 336, 321, and 345 residues were modeled for subunits A, B, and C, respectively, with Ramachandran outliers of 0.21%. These individual subunits have highly similar overall folding with global root-mean-square deviations (RMSDs) of 0.50 Å between subunits A and B, 0.64 Å between A and C, and 0.61 Å between B and C, introducing subtle structural adjustments when assembled into a T=3 lattice. The Basic, S, and P1 domains adopt distinct structural motifs; notably, the Basic domain forms a long α-helix that is structurally characterized for the first time in this study. The S domain adopts a typical jelly-roll β-barrel fold composed of eight antiparallel β-strands, whereas the P1 domain forms a squashed β-barrel consisting of seven antiparallel β-strands.

**Figure 2.**
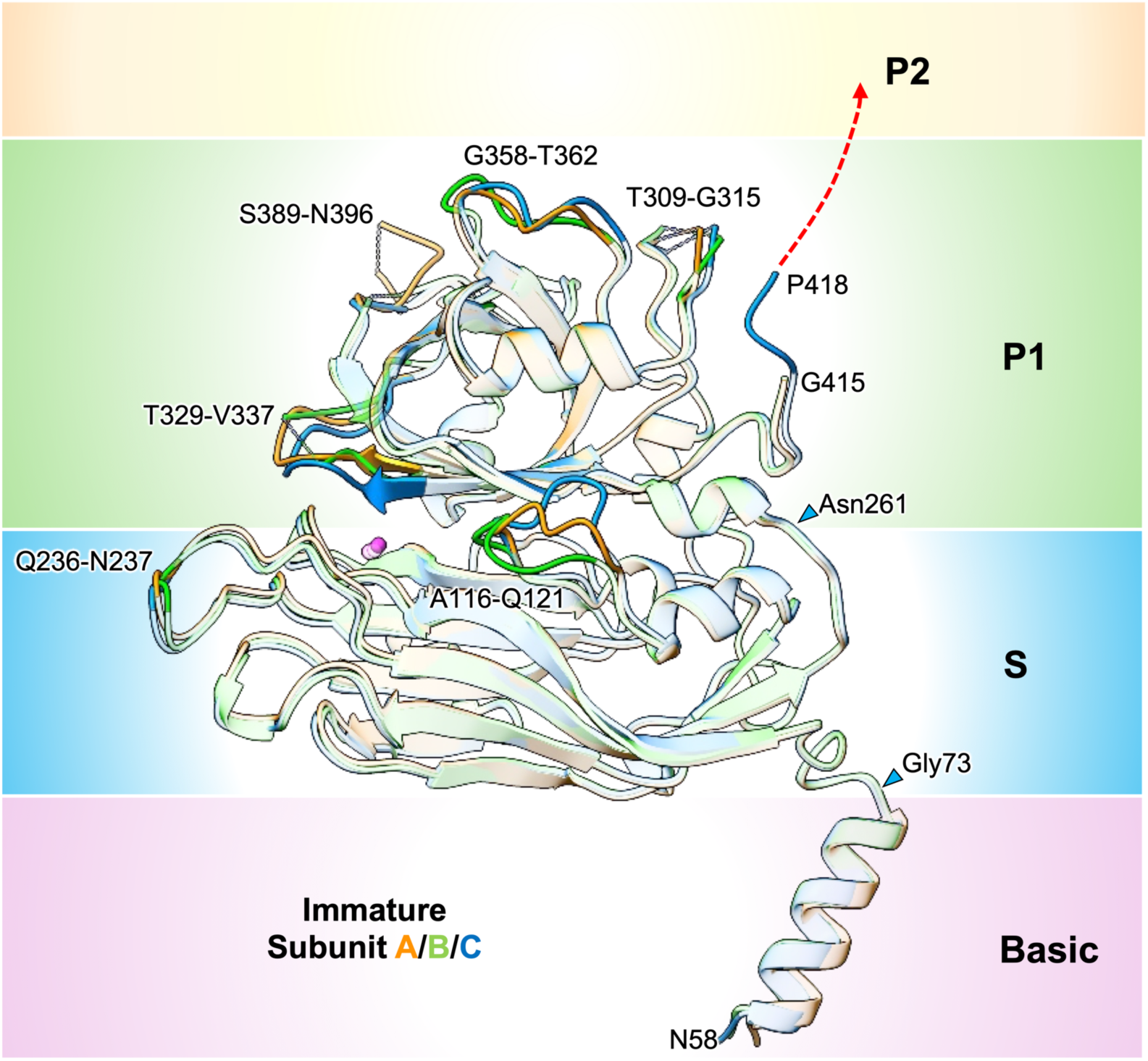
Structural comparison of capsid subunits A, B and C in the immature HAstV-4 particle. Atomic models of subunits A, B and C were superimposed to compare their structural differences within the capsid core. Ribbon representations are shown as transparent for regions with root-mean-square deviation (RMSD) <1.6 Å and opaque for regions with RMSD ≥1.6 Å. Regions of higher RMSD are coloured orange (subunit A), green (subunit B) and blue (subunit C). The capsid core domains are indicated as Basic, S and P1. Amino acid residues discussed in the main text are labelled.

### Subunit-specific proteolysis drives capsid maturation

The maturation of HAstV-4 is driven by the proteolysis of the capsid protein precursor VP70 into VP34 and VP27/VP25 (Fig. 1a). As maturation progresses, the VP27 fragment containing the P2 domain remains associated in the C-C dimers but is selectively removed from the A-B dimers.

Comparing the atomic models of subunits A, B, and C between immature and mature particles revealed trypsin-induced structural changes specific to each subunit (Fig. 3 and Supplementary Fig. 2).

**Figure 3.**
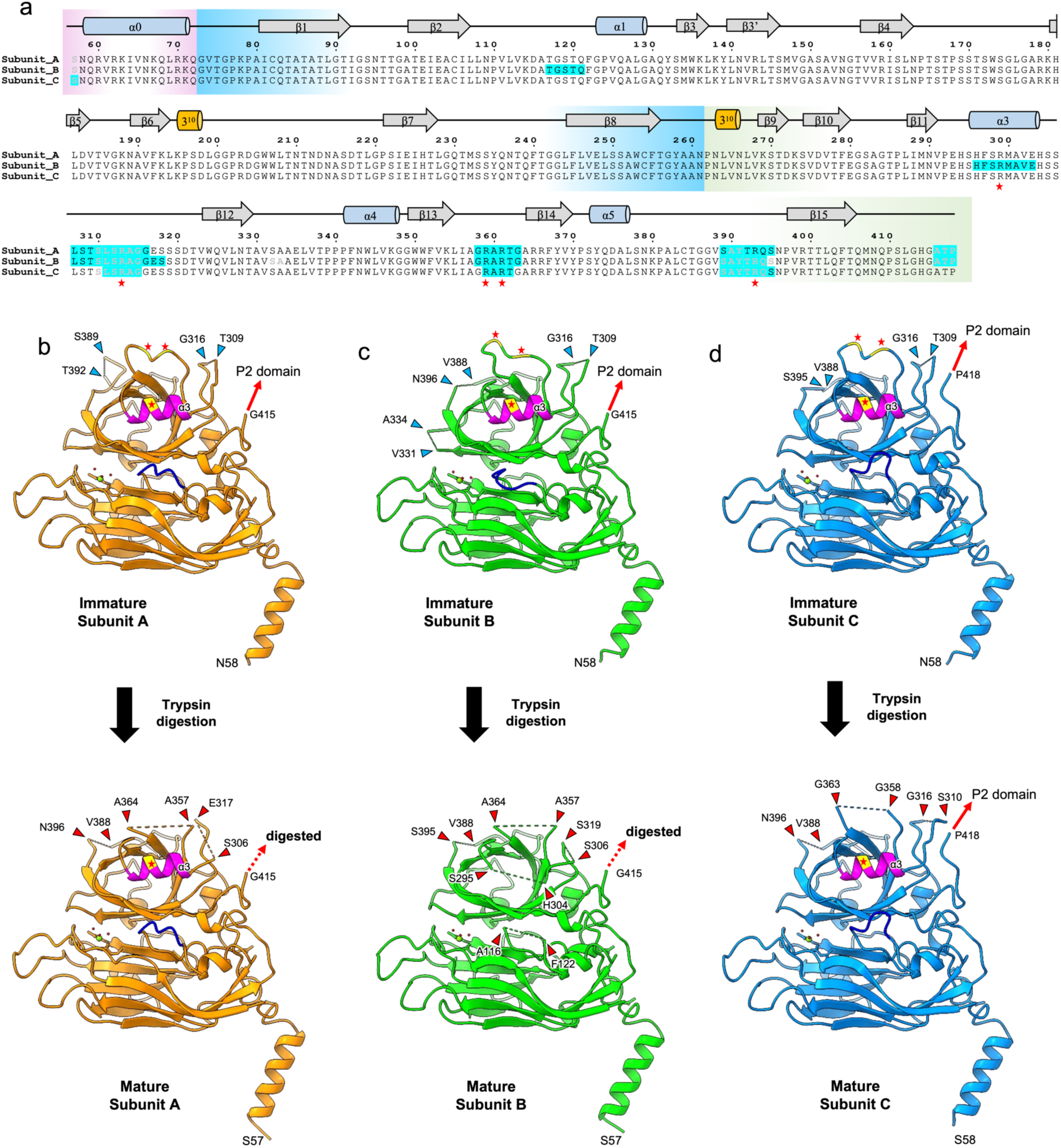
Subunit-specific structural changes in HAstV-4 capsid cores upon proteolytic maturation. (a) Amino acid sequences and secondary structure assignments of the capsid core for subunits A, B and C. Residues lacking density in immature and mature particles are indicated in grey and blue, respectively. (b-d) Comparison of subunits A (b), B (c) and C (d) in immature (top) and mature (bottom) particles. Structural models are shown to highlight regions that become disordered upon maturation. Secondary structure assignments were generated using ESPript (v3.2) ^36^ and further refined based on DSSP analysis in UCSF ChimeraX.

The most extensive structural changes occur in subunit B, where the loops between Ala116-Phe122 and between Ala357-Ala364 become disordered during maturation. The Ala116-Phe122 loop in the S domain originally exhibits distinct conformations in each subunit. It remains unaltered in subunits A and C but becomes disordered in subunit B after trypsin digestion. This is likely due to indirect structural changes because the loop lacks major cleavage sites. Conversely, the Ala357-Ala364 loop in the P1 domain originally shows identical conformations in subunits A and C but a different conformation in subunit B. This loop contains two major trypsin cleavage sites (Arg359 and Arg361) and becomes structurally disordered in all three subunits after digestion, confirming it as a direct target of proteolytic degradation.

The Ser295-His304 loop, including the short α3 helix, originally adopts the same conformation across all subunits. Despite containing the major trypsin cleavage site Arg299, this structure is maintained in subunits A and C during maturation, but completely disappears in subunit B, indicating that α3 helix digestion is modulated by the subunit-specific local environment. Furthermore, the originally disordered loops between Thr309-Gly316 and between Val388-Pro397, which both contain arginine cleavage sites, exhibit an increase in disordered residues after maturation, showing that their inherent flexibility is further enhanced by trypsin digestion.

In addition to local disordering, broader conformational changes occur between the S and P1 domains (Supplementary Fig. 3). In all subunits, a hydrated metal ion is stably bound to the S domain surface via Ser94 and Glu101. In immature subunits A and B, Thr329 of the P1 domain also interacts with this metal ion, maintaining S-P1 domain distances of 8.7 Å and 8.2 Å, respectively (Supplementary Fig. 3a,b). Upon maturation, the Thr329 side chain rotates away, reducing the domain distances to 6.5 Å. In subunit C, Thr329 does not originally contribute to metal coordination, and the S-P1 distance remains nearly unchanged (6.5 Å to 6.4 Å) (Supplementary Fig. 3c). The larger initial S-P1 domain distance mediated by the hydrated metal ion in subunits A and B likely facilitates their selective proteolysis, potentially leading to the release of the P2 domain.

### Distinct architectures of A-B and C-C dimers

Focused refinement yielded local reconstructions of flexible P2 domains at 5.35 Å for A-B dimers and 5.12 Å for C-C dimers in immature particles, and 4.75 Å for C-C dimers in mature particles (Fig. 4 and Supplementary Figs. 4 and 5). These local refinement maps revealed that the P2 dimers adopt distinct orientations and inclinations relative to the capsid surface between the mature and immature states (Fig. 4a and b). Rigid-body docking of a previously determined crystal structure (PDB 3QSQ) into these densities showed no significant direct inter-molecular interactions between the P1 and P2 domains in either A-B dimer or C-C dimer, suggesting that the P2 domains are primarily tethered via flexible linkers.

**Figure 4.**
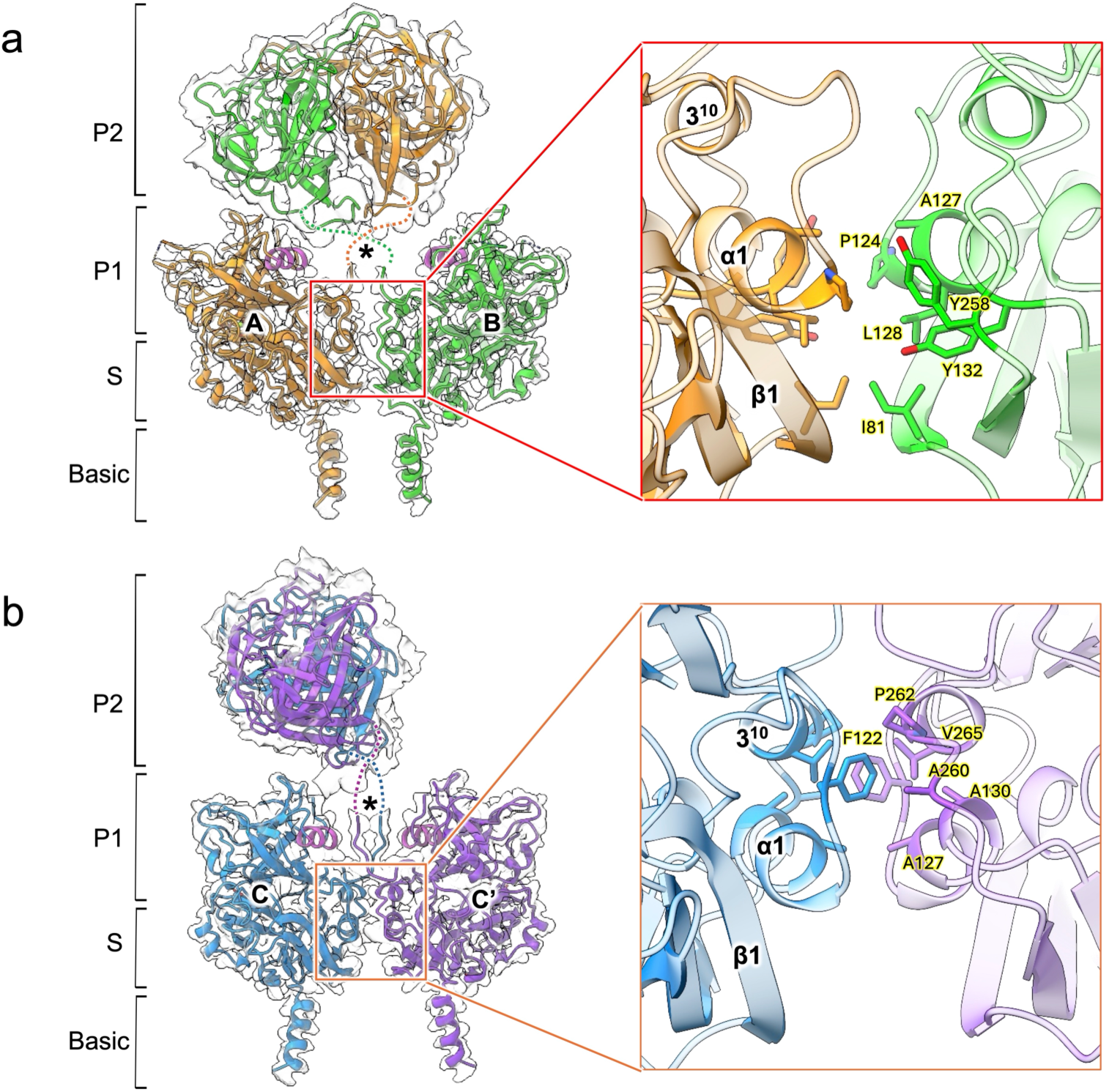
Comparison of intermolecular interactions in A-B and C-C dimers of the immature HAstV-4 particle. (a) Ribbon representation of the A-B dimer with the cryo-EM map (left) and a close-up view of the interaction interface within the capsid core (right). Subunits A and B are coloured orange and green, respectively. A broad cavity formed at the interface is indicated by a black asterisk. (b) Ribbon representation of the C-C dimer with the cryo-EM map (left) and a close-up view of the interaction interface within the capsid core (right). The two subunits C are coloured blue and purple. A narrower cavity at the interface is indicated by a black asterisk. The P2 domains shown in both dimers are based on the crystal structure of the HAstV-8 P2 domain (PDB 3QSQ) and are fitted into the cryo-EM maps.

**Figure 5.**
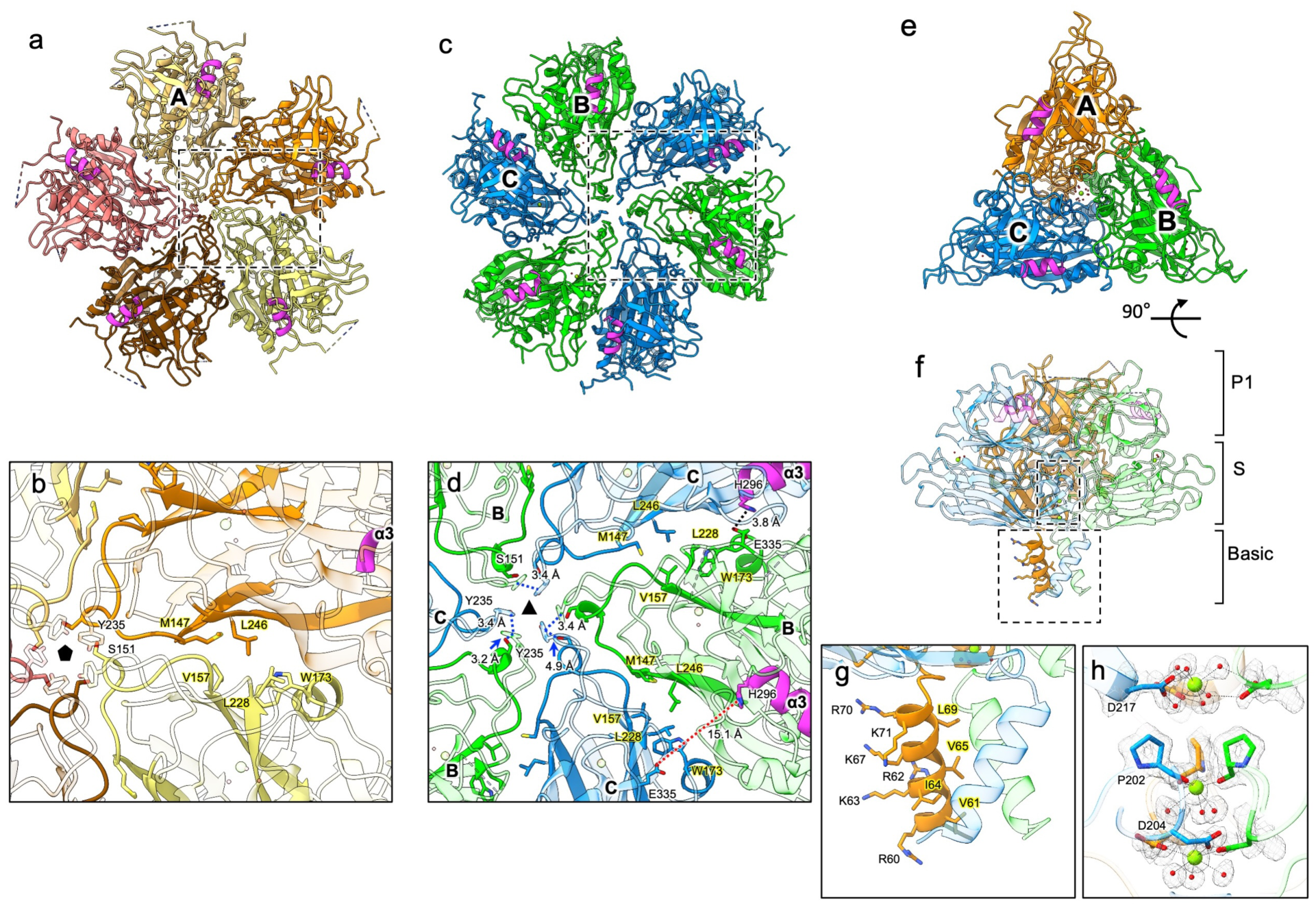
Intermolecular interactions within the HAstV-4 capsid core. (a) Intermolecular interactions of subunit A around the icosahedral 5-fold axis. (b) Close-up view of the boxed region in (a). Subunit A interacts with adjacent subunits through hydrophobic contacts, and the central axis is sealed by an array of Tyr235 side chains, supported by Ser151. (c) Organization of the icosahedral 3-fold axis formed by alternating subunits B (green) and C (blue). (d) Close-up view of the boxed region in (c). Subunit B interacts with neighboring subunit C molecules through hydrophobic contacts at one interface and π-π interactions between Tyr235 residues at the 3-fold axis. (e) Architecture of the pseudo-3-fold axis formed by subunits A (orange), B (green) and C (blue). (f) Side view of (e), showing the helical bundle formed by the N-terminal Basic domains. (g) Close-up view of the helical bundle, in which the three subunits are stabilized by hydrophobic interactions. (h) Densities consistent with coordinated metal ions are observed along the pseudo-3-fold axis.

When assembled into the capsid shell, these defined subunit blocks generate dimer-specific configurations with distinct overall architectures. In A-B dimers, subunits A and B interact primarily through the S domain, where the α1 helices and β1 sheets interact inversely with each other in an anti-parallel manner. This interface forms a hydrophobic core consisting of Ile81, Pro124, Ala127, Leu128, Tyr132, and Tyr258 residues (Fig. 4a). As a result of this assembly, a broad groove is formed on the shell surface directly underneath the P2 domain. In contrast, C-C dimers are stabilized by S and P1 domain interactions, where anti-parallel α1 and 3_10_ helices form a hydrophobic core consisting of Phe122, Ala127, Ala130, Ala260, Pro262, and Val265 residues (Fig. 4b). Consequently, a much narrower groove is formed underneath the P2 domain on the rigid shell surface of the C-C dimer compared to that of the A-B dimer. These differential dimeric interfaces demonstrate that dimer-specific architectures define distinct local environments on the capsid surface, directly influencing the spatial positioning, chemical accessibility, and structural variations in the localization of the P2 domain.

### Intermolecular interactions governing the T=3 icosahedral assembly

To understand the principles of HAstV-4 capsid assembly based on our atomic structures, we investigated the specific intermolecular interactions formed between adjacent subunits A, B, and C across the higher-order symmetry axes (Fig. 5). At the icosahedral 5-fold vertices, five subunit A monomers form a symmetrical homotypic interface stabilized by a hydrophobic core of Met147, Val157, Leu228 and Trp173 (Fig. 5a,b). The central strict 5-fold axis is filled with a hydrophobic core formed by Tyr235 residues extending from each subunit. These Tyr235 residues are further supported from the inside by surrounding loops containing Ser151, which provide essential structural reinforcement to stabilize the Tyr235 residue configuration.

Along the icosahedral 3-fold axes, subunits B and C assemble to form heterotypic interfaces, where adjacent Tyr235 residues engage in π-π stacking interactions at a distance of approximately 3.4 Å, effectively sealing the central axis (Fig. 5c,d). Notably, the structural support provided by Ser151 exhibits pronounced asymmetry between the two subunits. In subunit B, the distance between the hydroxyl group of Ser151 and the Try235 side chain is 3.2 Å. In subunit C, this distance increases to 4.9 Å, which no longer provides direct structural reinforcement. As a result, the Tyr235 residue is not structurally reinforced and becomes conformationally flexible. This unique configuration suggests that Tyr235 in subunit B is firmly fixed, while Tyr235 in subunit C flexibly seals the central 3-fold axis.

This structural asymmetry is enhanced by asymmetric intermolecular interactions between His296 and Glu355 (Fig. 5d). Glu355 in subunit B interacts with His296 in the adjacent right-hand subunit C at a distance of 3.8 Å, whereas His296 of subunit B is completely separated from Glu355 of the left-hand subunit C at a distance of 15.1 Å. Remarkably, this His296-Glu355 intermolecular interaction occurs exclusively between the specific subunit B and C pairs where no homotypic intermolecular interaction exists between the Tyr235 pair. These complex inter-subunit constraints and specific molecular configurations work together to precisely regulate the flexibility and structural heterogeneity of the local three-dimensional structures observed around the 3-fold symmetry axes.

Furthermore, local pseudo-3-fold axes are formed at the junctions of subunits A, B, and C (Fig. 5e). At this location, the N-terminal Basic domains of each subunit interact to form a prominent helical bundle extending directly into the interior of the virion (Fig. 5f,g). This bundle is stabilized by hydrophobic interactions among inwardly facing hydrophobic residues (Val61, Ile64, Val65, and Leu69), while positively charged residues (Arg60, Arg62, Lys63, Lys67, Arg70, and Lys71) face outward. Distinct densities corresponding to three metal ions were clearly identified along the pseudo-3-fold axis, where the metal ions are positioned between the S and P1 domains and precisely coordinated by the Asp204 side chains, the Pro202 main chains, and the Asp217 side chains via water molecules, respectively (Fig. 5h).

### RNA-associated features of the inner capsid surface

Cryo-EM maps revealed that the inner surface of the capsid shell is predominantly occupied by positively charged residues in both immature and mature particles (Fig. 6). This strong positive electrostatic potential is highly concentrated around the triple α-helix bundles formed by the Basic domains at the pseudo-3-fold symmetry axes (Fig. 6a). Notably, six unique and ordered unknown densities extend inward from Tyr139, repeating a disk-shaped pattern around these helical bundles (Fig. 6b). The spacing between these captured internal densities is approximately 3.3-3.7 Å, suggesting that this represents only a small subset of the 6.7 kb viral genome, and that these densities do not change during the maturation process.

**Figure 6.**
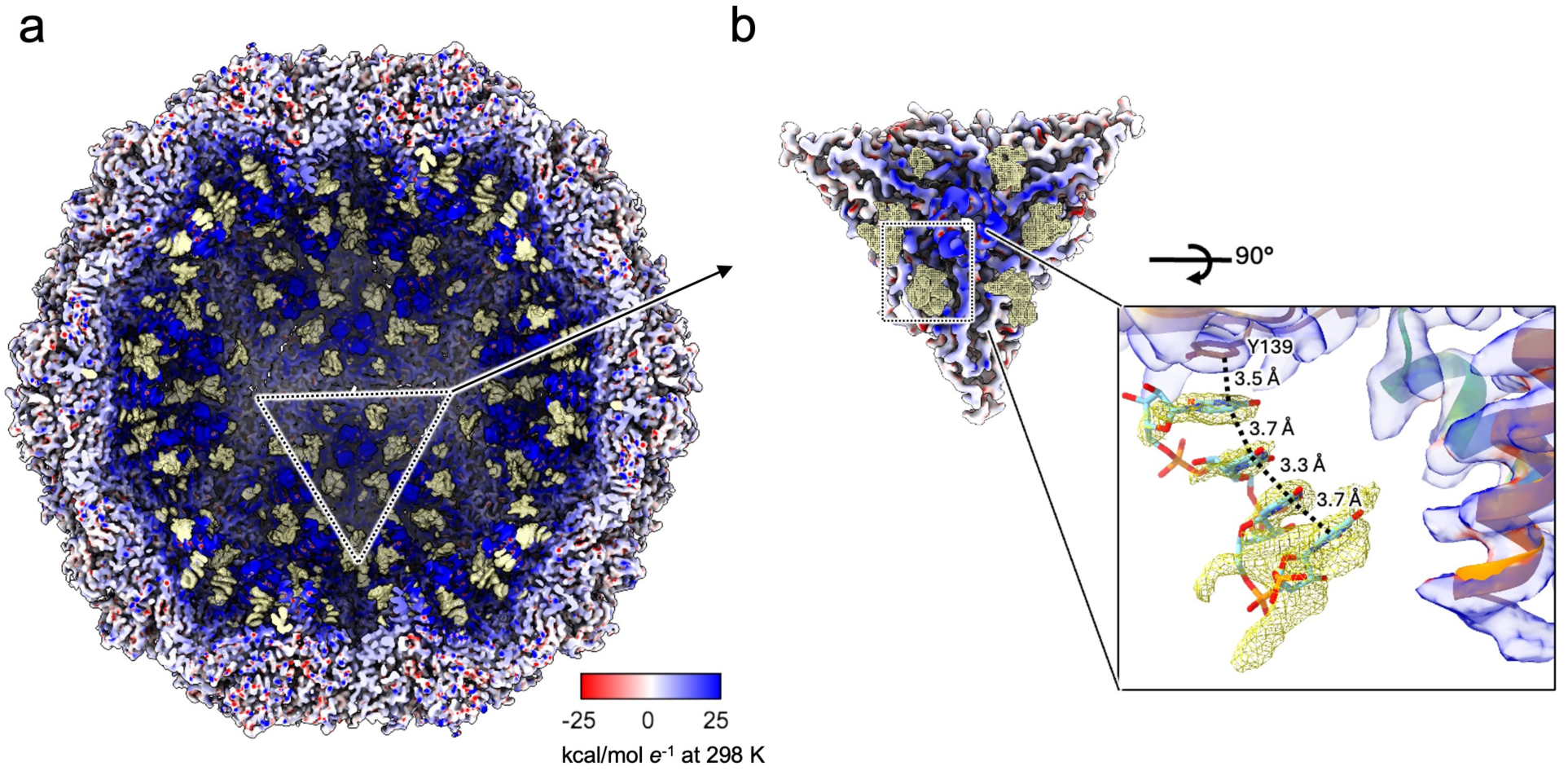
Electrostatic potential of the inner surface of the HAstV-4 capsid and putative interaction of viral RNA with the inner capsid surface. (a) The inner surface of the capsid is shown as a Coulombic electrostatic potential map, illustrating the distribution of positive (blue) and negative (red) charges using the ChimeraX coloured scheme. (b) Discrete internal densities observed near the pseudo-3-fold axes on the inner surface of the HAstV-4 capsid were tentatively fitted with a single-stranded RNA (ssRNA) model. In the close-up view, the model is consistent with the density and suggests potential interactions with residues in the S domain, including Tyr139. The electrostatic potential scale is expressed in kcal mol⁻¹ e⁻ at 298 K.

## Discussion

Comparison of our cryo-EM structures with previously reported crystallographic models of HAstV capsid proteins ^15,16^ revealed several important differences that emerged from the analysis of native particles at near-atomic resolution (Supplementary Fig. 6). In particular, loops at S-P1 interface, including the Ala116-Phe122 loop, which were unresolved or exhibited high B-factors in the crystal structures of HAstV-1 (PDB 5EWN) and HAstV-8 (PDB 5IBV), were clearly resolved in a subunit-specific manner (Fig. 3 and Supplementary Fig. 2). These observations reveal intrinsic conformational heterogeneity among quasi-equivalent subunits that could not be captured in previous X-ray crystallographic studies.

Our data demonstrates that capsid maturation is governed by subunit-specific proteolysis rather than a uniform process across the particle. Structural comparisons revealed that multiple loops become selectively disordered upon maturation, particularly the Ala116-Phe122 loop in subunit B. Notably, residues previously considered resistant to trypsin cleavage, such as Arg299 within the α3 helix, exhibit subunit-dependent behavior, indicating that local structural context influences proteolytic outcomes ^19^. These observations support a model in which capsid geometry determines protease accessibility. The three loops located at the P1 and P2 interface (Thr309-Gly316, Ala357-Ala364, and Pro389-Arg399) are highly conserved among HAstV-1, HAstV-4, and HAstV-8 and contain multiple trypsin cleavage sites, including Arg313, Arg359, Arg361, and Arg393 (Fig. 3a). Although these loops were well resolved in the crystal structures, Thr309-Gly316 and Pro389-Arg399 are intrinsically disordered in immature particles, and all three loops become disordered following trypsin digestion, as observed in cryo-EM maps.

Previous biochemical studies identified VP34, VP27 (beginning at Gln394), and VP25 (beginning at Ser424) as the major maturation products of HAstV-8 ^12–14^, establishing the classical VP34/VP27/VP25 maturation pathway. In this model, spike structures associated with VP27 remain attached to the particle, whereas VP25-containing fragments are released during maturation ^14,18^. Structural analysis suggests that the distinct geometries of the quasi-equivalent dimers differentially regulate access to these cleavage sites. In A-B dimers, a broad cavity approximately 30 Å in diameter is present beneath the P2 domain (Fig. 7a), providing proteases with access to multiple cleavage sites located at the P1-P2 boundary (residues 394-424). This open architecture may also permit cleavage at Arg299, resulting in the disruption of the α3 helix (residues Ser295-His304) specifically in subunit B (Fig. 3 and Supplementary Fig. 2), thereby facilitating subsequent processing to generate VP25. In contrast, C-C dimers contain a much narrower cavity, approximately 11 Å in diameter, beneath the P2 domain (Fig. 7b). This restricted geometry is likely to impose substantial steric constraints on protease entry, thereby protecting both the α3 helix containing Arg299 and the flexible P1-P2 boundary loops from proteolytic attack. Such protection may favor retention of VP27-associated spike structures on the particle surface. Together, these observations suggest that capsid geometry regulates not only overall susceptibility to proteolysis but also the extent of proteolytic processing within quasi-equivalent dimers. Differential accessibility of cleavage sites may therefore drive heterogeneous spike maturation and contribute to the selective release of VP25-associated fragments.

**Figure 7.**
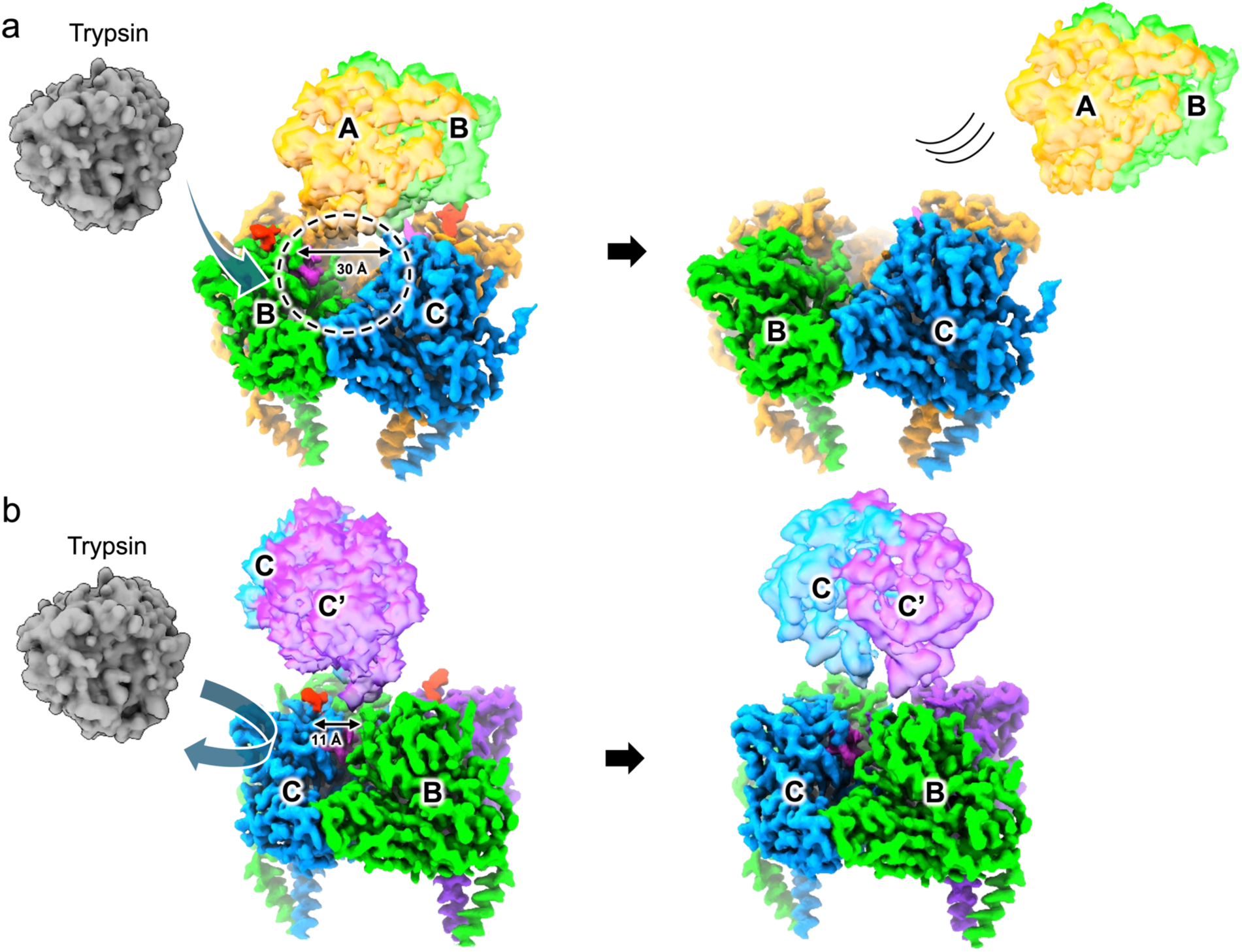
Structural basis for selective protease accessibility in A-B and C-C dimers. Cryo-EM maps of subunits A, B and C are coloured orange, green and blue (or purple), respectively. In the A-B dimer, a large cavity (∼30 Å in diameter) is formed between the P2 domain and the capsid core, permitting access for trypsin to reach cleavage sites. In contrast, the C-C dimer forms a much narrower cavity (∼11 Å), which restricts protease access. A model of trypsin is shown for reference and was generated in UCSF ChimeraX using the molmap function based on a previously determined structure (PDB 1TRN) ^37^.

In addition to localized proteolysis, cooperative rearrangements at the S-P1 domain interface associated with changes in metal-ion coordination were observed (Supplementary Fig. 3). The Val331-Glu335 loop, which was unresolved or highly flexible in previously reported crystal structures (Supplementary Fig. 6), adopts distinct subunit-specific conformations near Thr329 in immature particles (Figs. 2 and 3). Upon trypsin digestion, the Thr329 side chains in subunits A and B rotate and move away from the metal ion (Supplementary Fig. 3a,b), and the interface architecture of all three subunits converges to a common conformation resembling that of subunit C. The observation that the S-P1 gap distances in the metal-free crystal structures are 5.4 Å and 6.0 Å, respectively, which are shorter than those observed in our maps (Supplementary Fig. 6), suggests either that capsid assembly stabilizes metal binding or that metal-ion coordination contributes to capsid stability and modulates domain spacing during maturation ^20^. Although previous X-ray crystallographic studies of HAstV-1 and HAstV-8 suggested the presence of an intramolecular disulfide bond between Cys82 and Cys254 within the P1 domain, no such covalent linkage was observed in our cryo-EM maps (Supplementary Fig. 7). Instead, consistent with previous biochemical observations^20^, the density observed in this region is more consistent with a transiently coordinated magnesium ion, which may contribute to maintaining the structural integrity of the capsid core.

The identification of disk-shaped internal densities spaced at 3.3-3.7 Å around the pseudo-3-fold axes provides important insights into how the HAstV-4 capsid accommodates and organizes its single-stranded RNA (ssRNA) genome (Fig. 6). Although the local resolution is insufficient to define their atomic identity, their structural arrangement allows a theoretical model of single-stranded genomic RNA to be fitted into these densities. The observed spacing is consistent with genomic RNA configurations reported for tobacco mosaic virus (TMV) ^21^, supporting the hypothesis that these densities represent ordered RNA segments anchored by Tyr139 (Fig. 6b). The preservation of these internal genomic features in both immature and mature particles suggests that capsid maturation by proteolysis is uncoupled from genome release. Therefore, the subunit-specific proteolysis described here likely functions primarily to remodel the outer spike architecture for host-cell attachment and entry, whereas viral uncoating is triggered later during infection by distinct cellular cues encountered within the host cell.

Based on previous biochemical studies and the reported HAstV-1/FcRn complex structure (PDB 9DBT) ^22–26^, superimposition of the FcRn-bound structure onto our maps revealed that, in immature particles, FcRn can bind to the P2 domains of A-B dimers, whereas binding to C-C dimers is sterically restricted (Fig. 8). Furthermore, structural modeling suggests that five FcRn molecules can be accommodated around each fivefold axis, whereas only one molecule can be accommodated around each threefold axis, yielding a total of 80 potential FcRn-binding sites. In contrast, mature particles lacking A-B spikes are predicted to accommodate 60 FcRn molecules on the remaining C-C dimers. These predicted binding capacities are consistent with previous ELISA-based observations and support a model in which receptor engagement can occur in both immature and mature particles. Following receptor attachment and maturation, the newly exposed C-terminus generated at Arg299 may interact with the host-cell membrane and contribute to viral entry ^19^.

**Figure 8.**
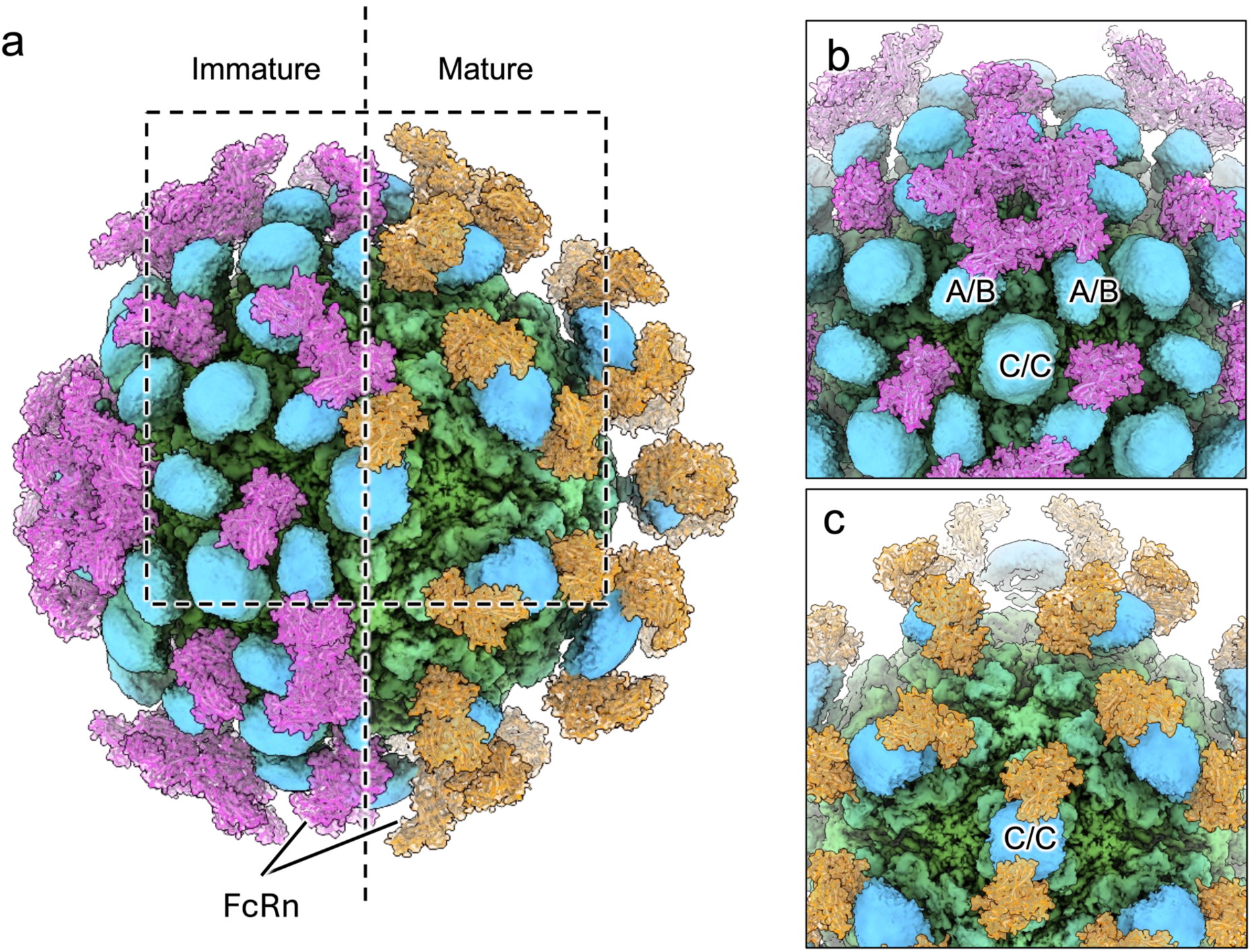
Structural modelling of FcRn binding to HAstV-4 particles. (a) Cryo-EM maps of mature (left) and immature (right) HAstV-4 particles with the P2-FcRn complex structure (PDB 9DBT) fitted into the maps. (b, c) Close-up views of the fivefold axis in the immature particle (b) and the threefold axis in the mature particle (c). P2 domains are coloured blue, and FcRn molecules are shown in purple (immature) or orange (mature). In immature particles, FcRn is predicted to bind primarily to A-B dimers, corresponding to a maximum of 80 binding sites. In mature particles, FcRn binding is restricted to C-C dimers, corresponding to 60 binding sites. The FcRn density was generated using the molmap function in UCSF ChimeraX.

Together, our near-atomic-resolution structures provide a comprehensive framework in which capsid architecture governs protease accessibility, enabling subunit-specific maturation that primes the virus for subsequent entry. By bridging the gap between crystallographic observations and the functional states of native viral particles, this study elucidates molecular mechanisms linking heterogeneous proteolysis, selective spike release, host-cell attachment, and viral uncoating. These findings establish a structural basis for proteolysis-driven activation in astroviruses and provide broader insights into maturation mechanisms employed by non-enveloped viruses.

## Material and Methods

### Preparation of immature and mature HAstV-4 particles

Caco-2/Cas9 cells were seeded into six T225 flasks and cultured to confluence. A frozen stock of HAstV-4 (GenBank accession number: LC930686) was activated with trypsin (type IX-S; Sigma-Aldrich, T0303-1G) at 10 μg ml⁻¹ and used to infect cells at a multiplicity of infection (MOI) of 1 for 1 h at 37 °C. After adsorption, cells were washed and maintained in minimum essential medium (MEM; Nacalai Tesque) for 2 days. Culture supernatants (180 ml) were clarified and layered onto a 30% (w/v) sucrose cushion, followed by centrifugation at 124,000 × g for 2 h (SW32 Ti rotor; Beckman Coulter). The pellet was resuspended in FBS-free MEM, adjusted to a density of 0.44 g ml⁻¹ with CsCl, and subjected to isopycnic ultracentrifugation at 150,000 × g for 24 h (SW55 Ti rotor; Beckman Coulter). Virus bands (∼1.35 g ml⁻¹) were collected, diluted tenfold in FBS-free MEM, and pelleted at 150,000 × g for 3 h. The final pellet was resuspended in 400 μl FBS-free MEM to yield immature particles. For preparation of mature particles, immature particles were treated with trypsin (type IX-S; 10 μg ml⁻¹), and the reaction was terminated by addition of FBS to a final concentration of 10% (v/v). The virus was then concentrated through a 30% (w/v) sucrose cushion at 124,000 × g for 2 h (SW32 Ti rotor), and the final pellet was resuspended in FBS-free MEM.

### Sample preparation for Cryo-EM

Quantifoil copper grids (R1.2/1.3, 200 mesh; Quantifoil Micro Tools) were glow-discharged using a plasma cleaner (PIB-10; Vacuum Device) prior to use. A 2.5 μl aliquot of immature or mature HAstV-4 particles was applied to the grids, which were then blotted at 4 °C and 100% humidity and plunge-frozen in liquid ethane using a Vitrobot Mark IV (Thermo Fisher Scientific). Grids were stored in liquid nitrogen until imaging.

### Cryo-EM data collection and image processing

Automated data acquisition was performed on a Titan Krios G4 cryo-electron microscope (Thermo Fisher Scientific) equipped with a cold field emission gun operated at 300 kV and a post-column Selectris X energy filter (10 eV slit width) (Table S1). Micrographs were recorded using a Falcon 4 direct electron detector in counting mode. Images were collected at a nominal magnification of ×130,000, corresponding to a calibrated pixel size of 0.92 Å, with a total electron dose of 40 e⁻ Å⁻² and a defocus range of −1.0 to −1.6 μm. Data were recorded in electron event representation (EER) format.

Image processing was performed using RELION (v3.1.4 and v5.0) (Supplementary Fig. 4 and 5) ^27,28^. Beam-induced motion correction was carried out using MotionCor2 ^29^, and contrast transfer function (CTF) parameters were estimated using CTFFIND (v4.1.14) ^30^. A total of 2,062 and 6,560 micrographs were used for immature and mature datasets, respectively, yielding final particle sets of 67,851 and 109,520 particles after classification and refinement. Initial 3D models were used as references for auto-refinement with imposed icosahedral (I1) symmetry. The resulting maps were subjected to masking and B-factor sharpening, followed by iterative rounds of CTF refinement and Bayesian polishing. Final maps were post-processed using a mask encompassing the entire particle. The global resolutions of the final reconstructions were 1.79 Å and 1.73 Å for immature and mature particles, respectively, based on the gold-standard Fourier shell correlation (FSC) criterion of 0.143. Local resolution estimation was performed using blocres in the Bsoft package ^31^.

### Focused refinement of P2 domains

For focused refinement of the P2 domains, symmetry expansion was performed in RELION using icosahedral (I1) symmetry. The P2 regions corresponding to A-B and C-C dimers were extracted from the expanded particles using ChimeraX ^32^ and RELION. This procedure yielded 55,448 and 50,728 particles for the A-B and C-C dimers of immature particles, respectively, and 589,877 particles for the C-C dimer of mature particles. Extracted particles were binned twofold, resulting in a pixel size of 1.84 Å, except for the mature dataset, which was processed without binning. Two-dimensional classification without image alignment was performed, followed by multiple rounds of three-dimensional classification with and without alignment, using a P2 domain reference map generated in ChimeraX. Final 3D auto-refinement was performed on the selected particle subsets. The resulting locally refined maps reached resolutions of 5.35 Å (A-B dimer, immature), 5.12 Å (C-C dimer, immature), and 4.75 Å (C-C dimer, mature), based on the gold-standard Fourier shell correlation (FSC) criterion of 0.143. Local resolution estimation was performed in RELION (v5.0).

### Model building and refinement

Initial atomic models were generated using ModelAngelo for both immature and mature particles ^33^. Cryo-EM maps corresponding to subunits A, B and C were extracted using UCSF ChimeraX. Models were manually adjusted and refined against the maps using Servalcat and Coot ^34,35^. Data collection, image processing and model validation statistics are summarized in Supplementary Tables 1, 2, and 3.

## Data availability

Cryo-EM maps of immature and mature HAstV-4 capsids, as well as focused reconstructions of the P2 domains, have been deposited in the Electron Microscopy Data Bank (EMDB) under accession numbers EMD-67811, EMD-67821, EMD-67822, EMD-67816 and EMD-67820. Atomic models have been deposited in the Protein Data Bank (PDB) under accession codes 21LZ and 21MB.

## Acknowledgements

This work was supported by the Japan Agency for Medical Research and Development (AMED) under Grant Numbers JP24fk0108667 and JP25fk0108667 (to K.K.), JP24fk0108669 and JP25fk0108669 (to K.M. and K.K.), and by the Research Support Project for Life Science and Drug Discovery (BINDS) from AMED under Grant Numbers JP24ama121005 and JP25ama121005 (to K.M.), NIPS Joint Research Program under Grant Numbers 25NIPS135 and 24NIPS118 (to K.K), ExCELLS Grant-in-Aid for Young Scientists Grant Numbers 24-Y4 and 25-Y13 (to K.H.).

## Author contributions

A.N. and K.K. provided virus strains. K.Mi., K.Ha., R.T.T. and K.K. purified virus particles. C.S. prepared cryo-EM grids and collected micrographs. K.Hi. and R.N.B.S. performed cryo-EM data processing. K.Hi. built atomic models. K.K. and K.Mu. conceived the study and acquired funding. K.Hi., K.K. and K.Mu. wrote the manuscript. All authors reviewed and revised the manuscript.

