## Supplemental Table 1-3 and Figure 1-7 for "Near-atomic structures of human astrovirus capsids reveal a subunit-specific maturation mechanism"

Running title: Subunit-specific maturation of human astrovirus capsids

**Supplementary Table 1. Cryo-EM data collection**

| **Data Collection** | |
|---|---|
| Electron microscopy | Titan Krios G4 |
| Detector | Falcon IV |
| Voltage | 300 kV |
| Nominal magnification | 130,000 |
| Total exposure time | 5.2 |
| Total electron dose | 40 electron/Å$^2$ |
| Number of EER fractionation | 25 |
| Nominal defocus range | -1.0 to -1.6 µm |

**Supplementary Table 2. Cryo-EM image processing**

| Image Processing | Immature particle | Immature particle P2 domain A-B dimer | Immature particle P2 domain C-C dimer | Mature particle | Mature particle P2 domain C-C dimer |
|---|---|---|---|---|---|
| Frame alignment | MotionCor2 | | | | |
| CTF estimation software | CTFFIND 4.1.14 | | | | |
| Number of micrographs | 2,062 | | | 6,560 | |
| 3D map reconstruction software | Relion v5.0 | | | Relion v3.1.4 and v5.0 | |
| Final Pixel size | 0.46 Å (super-resolution) | 1.84 Å | 1.84 Å | 0.46 Å (super-resolution) | 0.92 Å |
| Particles contributing to final map | 67,851 | 55,428 | 50,728 | 109,520 | 589,877 |
| Applied symmetry | I1 | C1 | C1 | I1 | C1 |
| Applied B-factor | -40 $Å^2$ | -314 $Å^2$ | -268 $Å^2$ | -40 $Å^2$ | -200 $Å^2$ |
| Global resolution (FSC = 0.143) | 1.79 Å | 5.35 Å | 5.12 Å | 1.73 Å | 4.75 Å |
| EMDB number | EMD-67811 | EMD-67821 | EMD-67822 | EMD-67816 | EMD-67820 |

**Supplementary Table 3. Validation statistics of the atomic models**

| Model Building | Immature particle (trimer) | Mature particle (trimer) |
|---|---|---|
| Modeling software | ModelAngelo, Coot, Servalcat, Molprobity | |
| Number of residues built | 350 (A), 342 (B), 349 (C) | 336 (A), 321 (B), 345 (C) |
| RMSD (bonds) | 0.0086 | 0.0079 |
| RMSD (angles) | 1.40 | 1.33 |
| Ramachandran outliers | 0.78 % | 0.82 % |
| Rotamer outliers | 1.28 % | 1.81 % |
| Clash score, all atoms | 8.29 | 7.39 |
| PDB ID | 21LZ | 21MB |

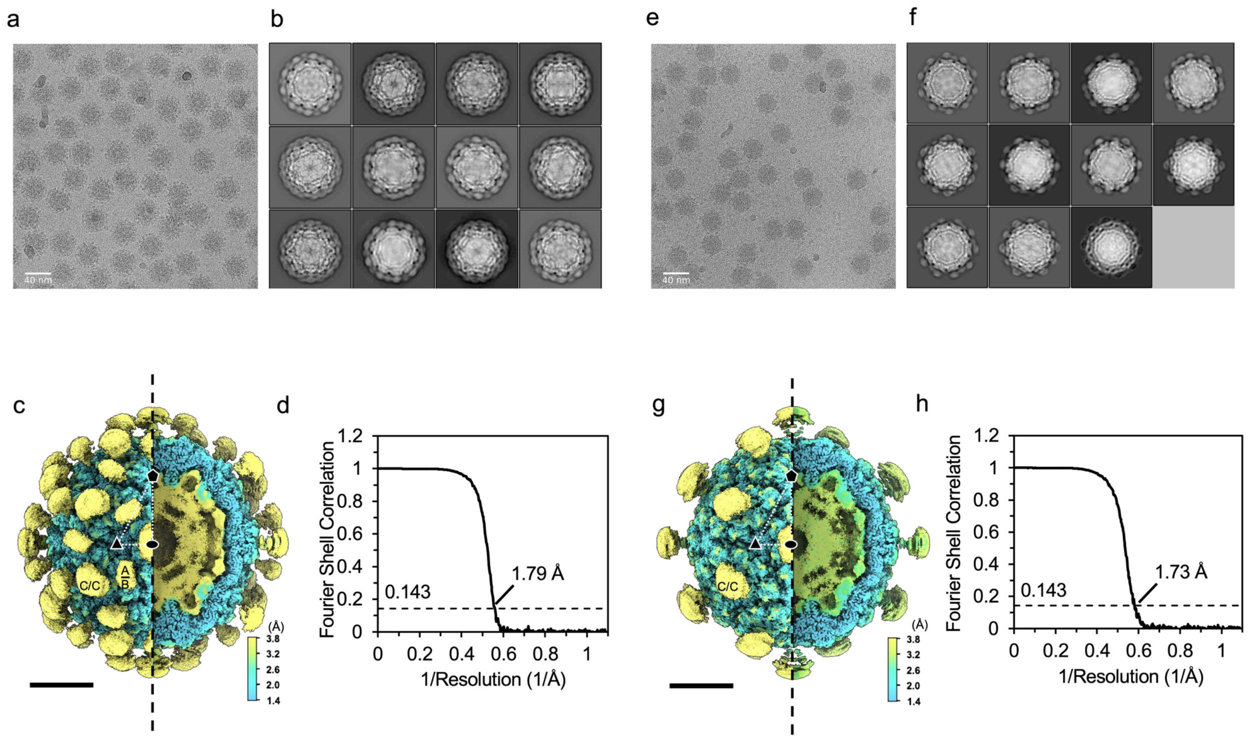


**Supplementary Fig. 1. Single-particle analysis of immature and mature HAstV-4 particles.** (a, e) Representative cryo-EM micrographs of immature (a) and mature (e) particles. (b, f) Two-dimensional class averages of immature (b) and mature (f) particles. (c, g) Local resolution maps of immature (c) and mature (g) particles, calculated using blocres in the Bsoft package. Surface views (left) and corresponding cutaway views (right) are coloured according to local resolution. Icosahedral twofold, threefold and fivefold symmetry axes are indicated by oval, triangle and pentagon symbols, respectively. The scale bar indicates 100 Å. (d, h) Gold-standard Fourier shell correlation (FSC) curves for immature (d) and mature (h) particles, with the 0.143 criterion indicated.

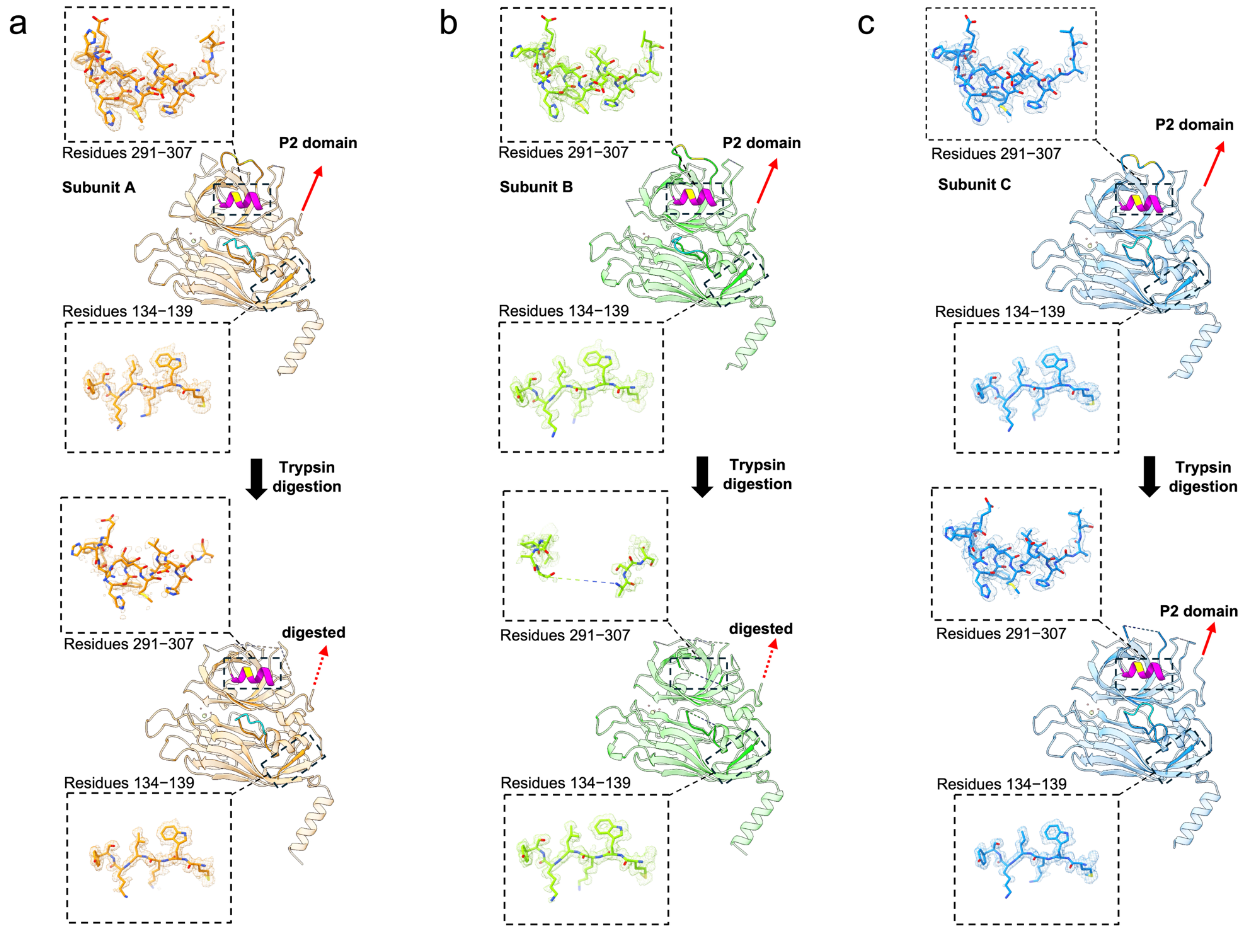


**Supplementary Fig. 2. Atomic models of immature and mature HAstV-4 subunit A, B, and C.** Ribbon diagrams of subunit A (a), subunit B (b), and subunit C (c) of immature (above) and mature (below), respectively. Cryo-EM density map of residues 134-139 (β3) and 291-307 (α3), coloured in magenta in ribbon model) superimposed with their atomic models.

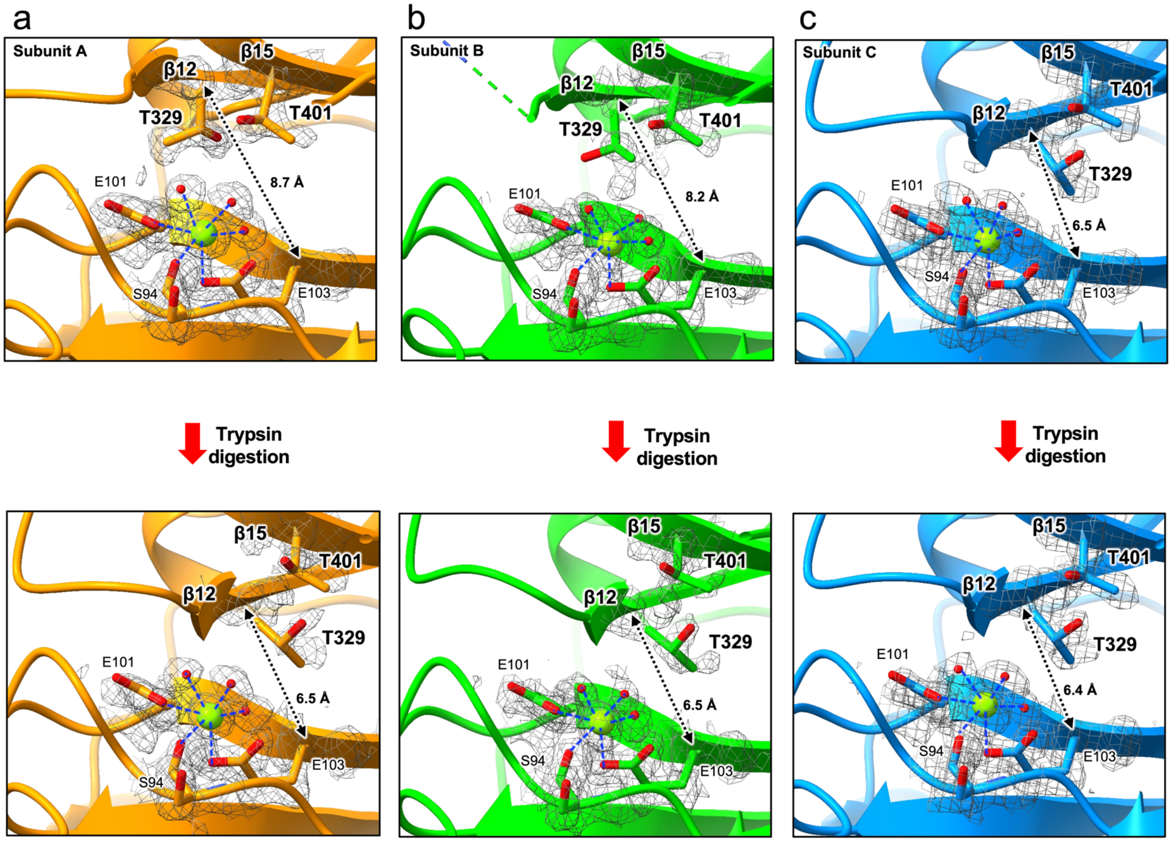


**Supplementary Fig. 3. Metal ion binding site before and after trypsin digestion.** Subunit A (a), subunit B (b), and subunit C (c) of immature (above) and mature (below), respectively. Water molecules are shown in red sphere and the metal ions are shown in green sphere. Interactions between the metal ion and surrounding atoms within 2.8 Å are indicated by blue dashed lines. The distances between Cα of Glu103 in the S domain and Cα of Thr329 in the P1 domain.

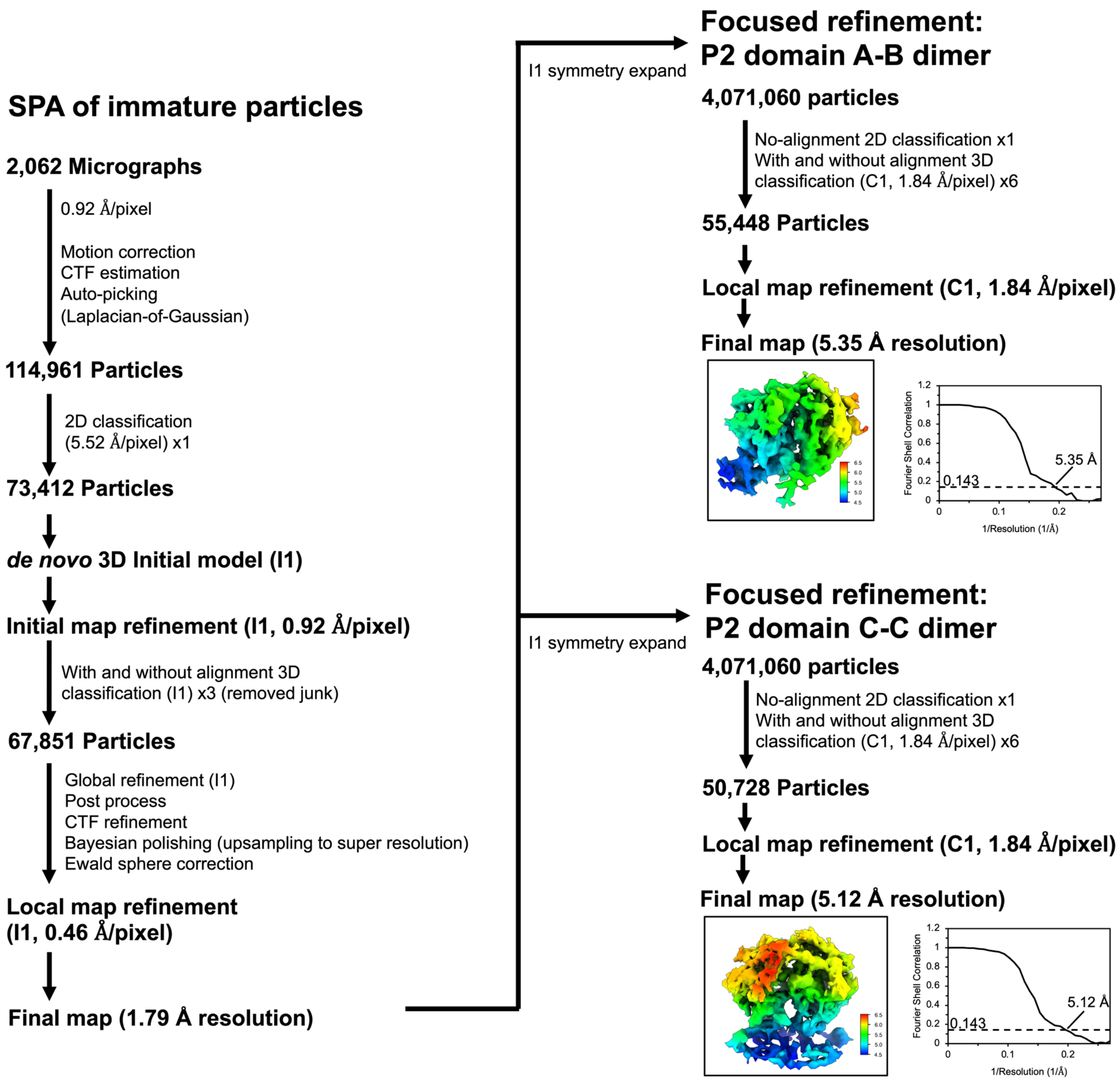


**Supplementary Fig. 4. Cryo-EM image processing workflow for immature HAstV-4 particles.** Flowchart illustrating the cryo-EM data processing and structure determination of immature HAstV-4 particles, including focused refinement of the P2 domains in A–B and C–C dimers.

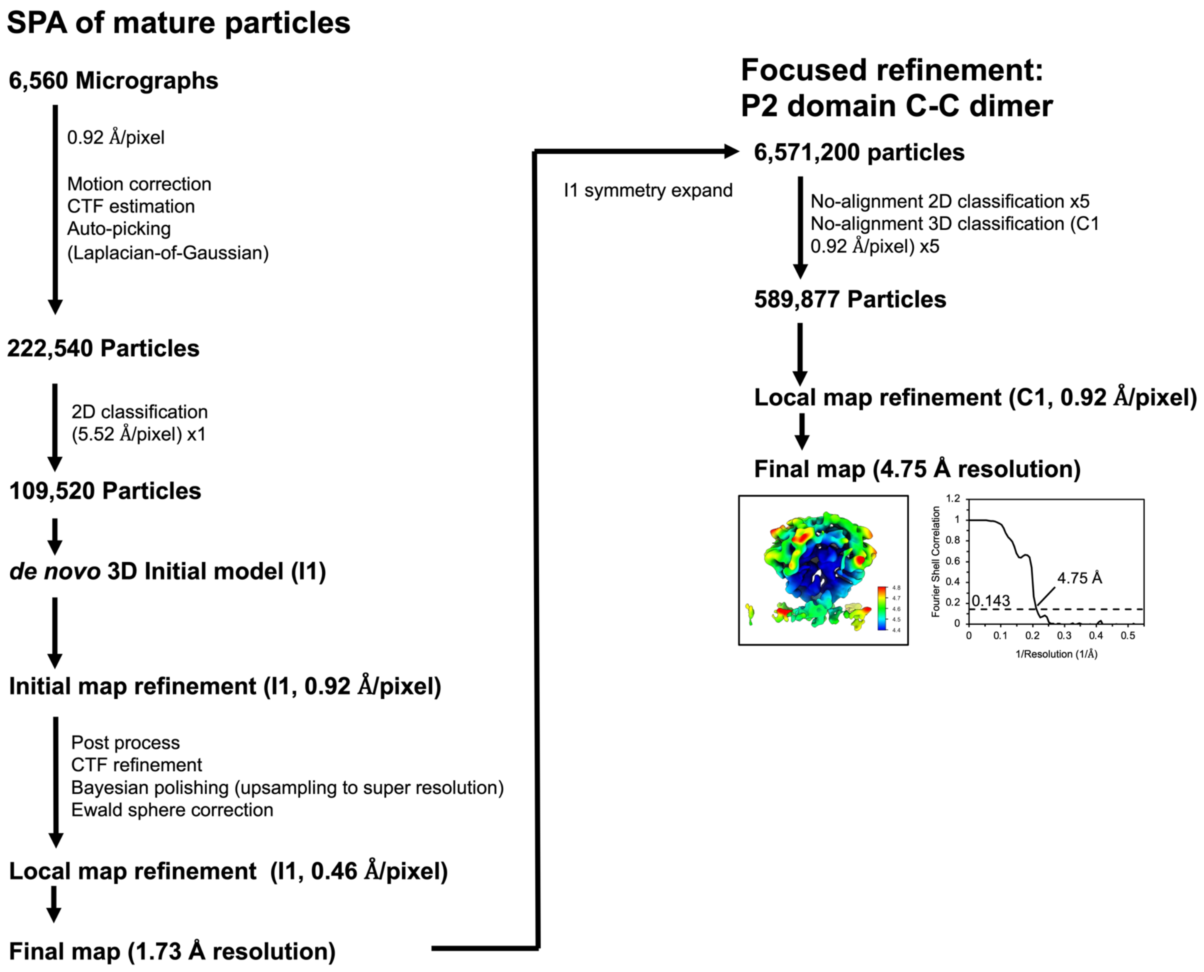


**Supplementary Fig. 5. Cryo-EM image processing workflow for mature HAstV-4 particles.** Flowchart illustrating the cryo-EM data processing and structure determination of mature HAstV-4 particles, including focused refinement of the P2 domains in C–C dimers.

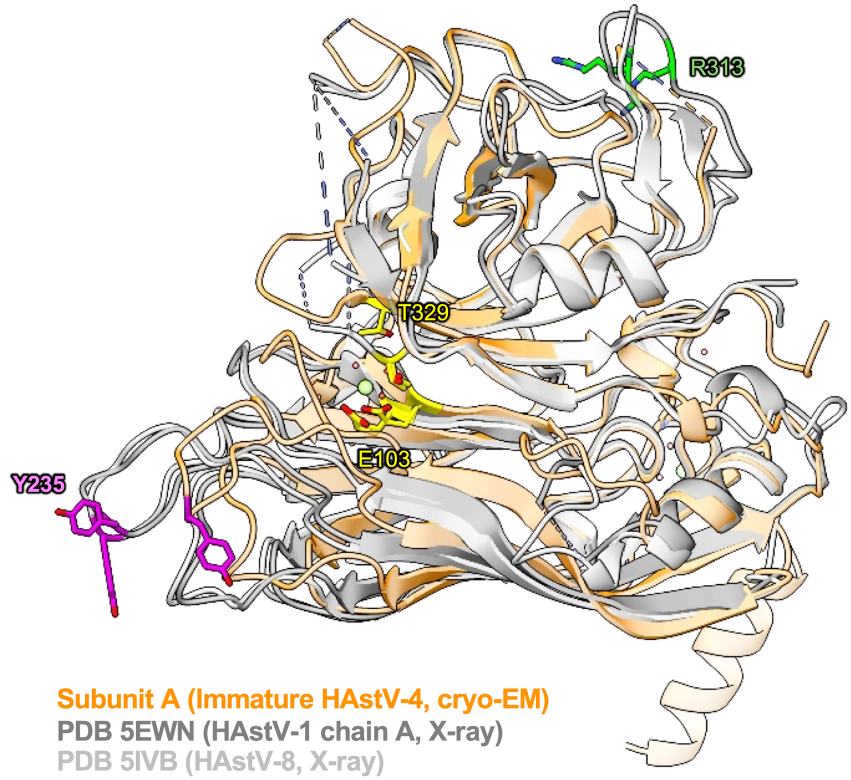


**Supplementary Fig. 6. Structural comparison between the HAstV-4 cryo-EM model and previously reported crystallographic models of the HAstV capsid proteins.** Superposition of the immature HAstV-4 capsid subunit A determined by cryo-EM (orange) with previously reported crystal structures of HAstV-1 VP34 (dark gray; PDB 5EWN) and HAstV-8 VP34 (light gray; PDB 5IVB). Regions exhibiting structural differences at the S–P1 interface are indicated.

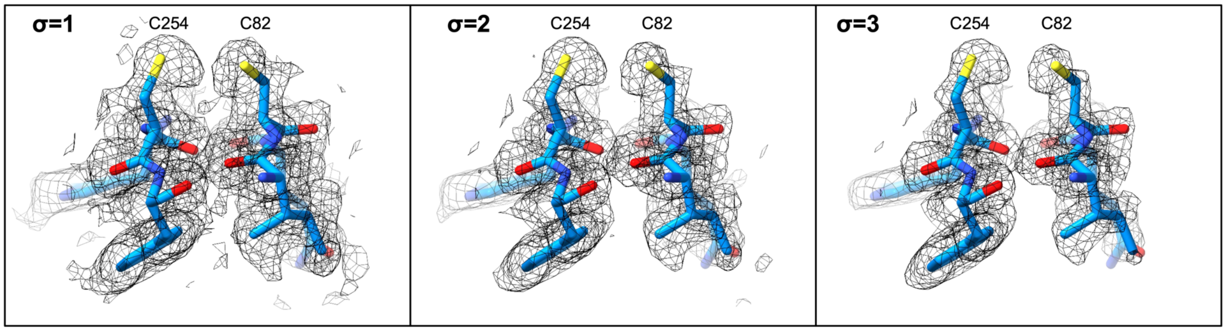


**Supplementary Fig. 7. Cryo-EM maps around Cys82 and Cys254 residues at different sigma levels.** The disulfide bond density between Cys82-Cys254, previously reported by X-ray crystallography[15,16], is not observed in the cryo-EM maps. The model of subunit C from the immature particle is shown as a representative example.
